# Characterisation of CD4 Th subsets as well as dietary, exercise and lifestyle factors in an established Rheumatoid Arthritis cohort; pilot study

**DOI:** 10.1101/2025.02.16.638543

**Authors:** Shweta Venkataraman, Nathan D’Cunha, Maria Kozlovskaia, Nenad Naumovski, Andrew McKune, Anne Bruestle, Tony Xu, Julien C Marie, Hector Hernandez-Vargas, Anna Dorai Raj, Ted Tsai, David B. Pyne, Chloe Goldsmith

## Abstract

**Background:** Sex disparities in the incidence and severity of Rheumatoid Arthritis (RA) are well-documented, with females experiencing more aggressive disease and different responses to treatment. While underlying mechanisms remain elusive, immune factors and lifestyle components, such as diet and physical activity, could play a role in shaping disease outcomes. Since Th cells are key drivers of RA, this study characterised CD4 Th subset frequencies, including Th1, Th2, Th17 and Th1/17 cells, in a cohort of males and females with established RA. Considering the role of diet and lifestyle in RA development, we also evaluated participant diet and lifestyle patterns.

**Methods:** An observational cross-sectional cohort study profiling 39 individuals (19 RA and 20 age and sex matched controls) with no major co-morbidities was conducted. Percentages of Th1, Th2, Th17 and Th1/17 cells from peripheral blood were determined by flow cytometry as well as serum levels of RF and cytokines IL17α, IFN-γ, TNF-α, GM-CSF, TGF-β, IL-6, and CRP by ELISA. Dietary intake was assessed by Food Frequency Questionnaire (FFQ), the Dietary inflammatory Index (DII) calculated from nutrient intake. Physical activity levels were evaluated by Global Physical Activity Questionnaire (GPAQ).

**Results:** Sex-specific differences of Th17, Th1 an Th2 cells were observed in RA groups. Males had lower levels of Th1 cells and higher levels of Th17 cells than control subjects, while females had lower levels of Th2 cells than controls. Female RA subjects had higher ratio of Th1:Th2 cells than controls, while male RA subjects had higher Th17:Th1 ratio than controls. Interestingly, circulating CD4 Th subset levels were predictive of RA in this cohort. No changes in cytokine levels were observed. The male RA group consumed 30% less carbohydrate (RA = 156g +/-57g, Control = 253g +/-59g; *p=*0.003), 37% less sugar (RA = 74.1g +/-29g, Control = 118g +/-33g; *p=*0.004) and 30% less dietary fibre (R = 33g +/-8g, Control = 48g +/-13 g; *p=*0.008) than the male control group. The female RA group had the lowest consumption of alcohol in comparison to female control group (RA = 0.31g, Control = 8.97g; *p*=0.047). Male subjects had a 4x higher DII than females (*p=*0.012).

**Conclusions:** Our findings highlight potential sex-specific differences in CD4+ Th cell subsets in individuals with established RA, indicating that sex may influence immune cell composition in this disease. Furthermore, dietary components showed sex-specific consumption patterns, underscoring the potential role of tailored lifestyle interventions in modulating immune responses in RA. Evaluating immunological sex differences and dietary and lifestyle factors may be important for enhancing management strategies in RA.

## Introduction

Rheumatoid arthritis (RA) is a chronic autoimmune polyarthritis with an estimated burden of up to 1% of the global population^1^. RA is characterised by synovial inflammation, vasculopathy, cartilage erosion and bone resorption^2^. The occurrence of disease and progressive joint destruction can cause drastic deterioration in functional ability and quality of life^3^. In RA, sex disparities are evident in treatment response and adverse effects, indicating potential underlying immunopathological and mechanistic differences^4^. Despite this, non-discriminatory recommendations between sexes in RA treatment is the current standard of care. Moreover, sex and gender discrepancies are still under-reported in clinical trials.

Although the precise aetiology of RA remains elusive, the interplay of environmental and genetic factors can trigger an adaptive autoimmune response long before clinical onset^5,6^. The involvement of CD4 T helper (Th) cells in RA pathology has been established^7,8^. In RA, CD4 Th cells constitute a large proportion of infiltrated inflammatory cells in the inflamed synovial membrane. CD4 Th cells form a complex network with other immune cells that promotes the pro-inflammatory cytokines, which trigger the activation of resident fibroblast-like synovial cells leading to cartilage and bone damage^9^. Specifically, an overactive Th1 pro-inflammatory, and overwhelmed Th2 regulatory response is postulated^9^. CD4 Th cells, and particularly Th17 cells, are endowed with a high degree of plasticity, defined as the ability of a cell to adopt characteristics of another Th cell subset. Th17 cells can acquire a Th1 cell profile expressing both IL-17 and IFNγ, referred to as Th1/17 cells^8,10,11^. This cellular population is thought to be highly pathogenic in RA and contribute to bone erosion by amplifying TNF and IL-1 expression^8^. However, the factors that drive Th1/17 plasticity in RA are not well understood. Characterisation of Th populations in RA at different stages of disease is warranted to shed light on the mechanisms underpinning their activity throughout the life of disease.

The management of RA typically involves life-long pharmaceutical intervention^12^. However, side effect burden can be considerable, and many individuals fail to reach remission. Therefore, it is common to look to modifiable lifestyle factors, such as diet, to aid in management^13^. Adherence to an anti-inflammatory diet, especially a Mediterranean Diet (MD), can improve symptoms and clinical symptoms of RA^14,15,16,17^, while specific dietary components can reduce risk of RA onset^18,14^ and others are currently prescribed as effective treatments for mild RA, such as fish oil containing omega 3 supplements. These studies support the proposition that dietary components have an intrinsic link to RA pathological mechanisms. While diet is an environmental factor that impacts virtually all aspects of host physiology^19^, the influence of dietary components on the immune system are only starting to emerge^20^. Indeed, certain dietary components can modulate the inflammatory profile of CD4 Th cells^21,11^. However, few immunological studies characterise lifestyle variables such as diet and physical activity in patients. Therefore, it is important to understand relationships between diet and disease, but also to provide effective dietary advice for individuals with RA.

In this pilot study we characterised the diet, lifestyle, cytokine levels and CD4 Th cell profiles in individuals with RA and age-sex matched healthy controls. This work reveals that Th subsets exhibit sex-specific profiles in established RA with negligible effects on cytokine expression. Dietary consumption patterns revealed certain factors may play a role in sex-specific immunological profiles, but more work is needed to validate these findings in larger cohorts.

## Methods and Materials

### Participants

An observational cross-sectional cohort study was conducted with 39 individuals (19 with RA (9 male and 10 female) and 20 age- and sex-matched controls). The inclusion criteria for the RA group included a formal diagnosis of RA from a Rheumatologist according to usual classification criteria at the time of diagnosis, in the absence of any other chronic conditions. Rheumatoid participants were mostly taking hydroxychloroquine and methotrexate, several patients were also on biological disease modifying antirheumatic drugs (DMARDs) and 5 participants were taking prednisone at the time of the study (Supplementary table S2). Participants in the control group were free of infectious diseases, malignant diseases, cardiovascular complaints, or chronic conditions. Participants were recruited through local and online media, rheumatologist referral and local community group newsletters including Arthritis Australian Capital Territory (Arthritis ACT). This project was approved by the University of Canberra Research Ethics Committee (approval number 20229197). Participants were informed of the study aims, benefits, risks and procedures, and they provided written informed consent before commencement.

### Characterisation of cohort

Participants completed one clinic visit at the University of Canberra. During their visit, we collected demographic, socioeconomic, and health-related information were collected by questionnaire from all participants. Anthropometric measurements, including participant body mass and height were collected for calculation of BMI according to World Health Organisation (WHO) standards^22^. The health-related quality of life of individuals with RA was assessed with the short form of Arthritis Impact Measurement score (AIMS2-SF) questionnaire^23,24^. Capability of individuals to regulate eating habits were evaluated using the Self-Regulation of Eating Behaviour Questionnaire (SREBQ)^25^. Risk assessment of hazardous alcohol intake was performed with the Alcohol Use Disorders Identification Test-Concise (AUDIT-C)^26,27^. The physical activity levels of this cohort were determined using the Global Physical Activity Questionnaire (GPAQ) developed by the WHO^28^.

### Determination of dietary component intake

An estimation of the daily energy (kcal) and nutrient intake (g/mg/μg) of all participants was obtained via an interview-administered validated food frequency questionnaire (FFQ)^29,30^, adapted from the Commonwealth Scientific Industrial Research Organisation (CSIRO)^31^. The FFQ contains 225 food items, covering all food groups and account for episodic consumption of food items including supplements. The FFQ data was analysed using FoodWorks^TM^ v10.0.4266 (Xyris Software, QLD, Australia) and the Australian Food Composition Database (Ausfoods 2017).

### Diet Inflammatory Index (DII)

The DII was calculated following the established methodology with some modifications^32^. In this study, we included 38 energy-adjusted food parameters. Scores for this index range from -8.87 (strongly anti-inflammatory) to 7.98 (strongly pro-inflammatory).

### Blood processing

A 40 mL sample of peripheral blood was collected by venepuncture in ethylenediamine tetraacetic acid (EDTA) tubes. Peripheral blood mononuclear cells (PBMCs) and plasma were separated by density gradient centrifugation at 1200 g for 10 min using a densitometry reagent (lymphoprep) and SepMate^TM^ tubes (StemCell Technologies, USA). PBMCs were isolated and washed twice with 2% fetal bovine serum (FBS) after centrifugation at 300 g for 8 min. Cell viability was determined by trypan blue staining and counting with Countess^TM^ 3 cell counter (Invitrogen, USA). PBMCs were labelled with a series of fluorochrome-conjugated antibodies (Supplementary Table 1) against surface markers according to manufacturer instructions (BioLegend, USA). After 30 min incubation in the dark at 4°C, all samples were washed with 2% FBS and centrifuged at 300 g for 5 min at 4°C. Cell pellets were resuspended in 2% FBS. Samples were analysed by FACSAria Fusion flow cytometer (Becton, Dickinson & Company, USA) within 4 hours of blood collection.

### Flow Cytometry

Cells were gated based on forward and side scatter characteristics and isotype controls for each specific marker. To avoid false positive PE-results and for compensation in multi-colour analysis, instrument calibration was conducted each day the samples were run. The frequency of Th cell subsets (Th1 (CXCR3+CCR4-CCR6-), Th2 (CXCR3-CCR4+CCR6-), Th17 (CXCR3-CCR4+CCR6+) and Th1/17 (CXCR3+CCR4-CCR6+)) were calculated using FlowJo^TM^ software v10.8.1 (Becton, Dickinson & Company, USA) as previously described ^33^. The gating strategy is presented in Supplementary Figure 1.

### Enzyme-linked immunosorbent assay (ELISA)

Cytokines from plasma were analysed using plate-based ELISA-kits according to manufacturer’s instructions. Interleukin-17 (IL17a) (88-7176-22, Thermo Fisher Scientific), granulocyte-macrophage colony-stimulating factor (GMCSF) (88-8337-22, Thermo Fisher Scientific), interferon-gamma (IFNγ) (88-7316-22, Thermo Fisher Scientific), transforming growth factor beta (TGFβ) (88-50390-22, Thermo Fisher Scientific), Tumour Necrosis Factor alpha (TNFα) (88-7346-22, Thermo Fisher Scientific), rheumatoid factor (RF) (RFAD11-50, Alpha Diagnostic) and C-reactive protein (CRP) (KHA0031, Thermo Fisher Scientific).

### Statistical Analysis

The distribution of data was tested with the Shapiro-Wilk test of normality. Descriptive statistics for normally distributed continuous valuables are presented as mean ± SD, while non-normally distributed data are presented as median (1^st^, 3^rd^ quartile). Categorical data are presented as frequencies and were tested for associations with the Fisher’s exact test. Th cell subsets are presented as percentage of total CD4 Th cells calculated as percent of parent (CD4+/CD8-). Unpaired two tailed Mann-Whitney tests were performed to compare Th subset frequencies.

A PacMAP was run on default parameters on all CD4 pre-gated T cells with no downsampling. Features were generated by manual identification of populations gated on the PacMAP and manually back-gated to identify and verify populations. Highest feature importance derived features from XGBoost classification on patient-nominated RA state, with feature importance’s accumulated across 10 iterations, each iteration consisting of a KFold repetition (splits = 3). The top 5 features were then selected, which coincided with the value that yielded the highest adjusted R, indicating that further inclusion of variables did not improve the model beyond chance. Subsequently, each of the lifestyle variables was correlated to each of the identified immune factors using a linear model, alongside the participant’s age, sex, BMI, and smoking level. Lifestyle variables and their sex interaction’s contribution towards the model were recorded and adjusted subsequently using a two-stage false discovery rate adjustment. Remaining statistically significant lifestyle variables have been presented. Statistical software R studio and GraphPad Prism (GraphPad Software, USA) was used for data analysis and graph preparation. P-values lower than 0.05 were considered statistically significant.

## Results

### Cohort characteristics

A total of 39 participants enrolled in the study. The RA group included a total of 19 participants including 10 females (mean age: 60.0 ± 9.3 yrs) and 9 males (mean age: 57.8 ± 12.2 yrs). The control group consisted of 20 participants, 10 females (mean age: 59.5 ± 11.7 yrs) and 10 males (mean age: 56.7 ± 15.1 yrs). There was no difference in the mean age of participants per group (*p=*0.918) **(Table 1)**. There was no difference in the AIMS2-SF mean symptom scores between the sexes in RA group (female score: 4.22 ± 1.10, male score: 4.81 ± 0.98, *p=*0.240), nor in the years since RA diagnosis (female: 11.0 ± 11.1, male: 15.3 ± 12.1. yrs, *p=*0.158). A large percentage of participants (70% in RA females and 88% of RA males) were taking disease-modifying anti-rheumatic drugs (DMARDs) and non-steroidal anti-inflammatory drugs (NSAIDs) (**Supplementary Table 2**).

**Table 1.** Socio-demographic information of participants in present study. AIMS2-SF: Arthritis Impact Measurement score short form. DMARDs: Disease modifying anti-rheumatic drugs, NSAIDs: Non-steroidal anti-inflammatory drugs. Data presented as mean ± standard deviation.

|  |  | Female subjects |  | Male subjects |  | p value |
| --- | --- | --- | --- | --- | --- | --- |
| Group |  | RA<br>(n=10) | Control<br>(n= 10) | RA<br>(n=9) | Control<br>(n= 10) |  |
| Age (years) |  | 60.0± 9.33 | 59.5 ± 11.7 | 57.4 ± 12.15 | 56.7± 15.1 | 0.9180 |
| BMI (kg/m<sup>2</sup>) |  | 28.8 ± 11.5 | 24.5 ± 3.89 | 27.9 ± 4.66 | 26.9 ± 4.32 | 0.5636 |
| AIMS2-SF score |  | 4.22 ± 1.10 | - | 4.81 ± 0.98 | - | 0.2395 |
| Time since RA diagnosis<br>(years) |  | 11.0 ± 11.1 | - | 15.3 ± 12.2 | - | 0.1575 |
| **Medication** |  |  |  |  |  |  |
| DMARDS, *n* |  | 7 | 0 | 8 | 0 |  |
| NSAIDS, *n* |  | 4 | 2 | 7 | 1 |  |
| RF, *n* |  | 6 | - | 4 | - |  |
| CRP (mg/dl) |  | 4.2 ± 3.8 | 6.5 ± 2.7 | 4.7 ±3.5 | 6.0 ± 2.9 |  |
| Physically Active, *n* |  | 9 | 7 | 8 | 9 | 0.9999 |
| Smoking history, *n* |  | 3 | 1 | 7 | 4 | 0.1837 |
| Current smoking, *n* |  | 0 | 0 | 0 | 0 |  |
| **Self-regulation of eating**<br>**habits** |  |  |  |  |  |  |
| Low |  | 0 | 1 | 2 | 0 |  |
| Medium |  | 6 | 4 | 2 | 3 |  |
| High |  | 4 | 4 | 5 | 6 |  |
| **Reported income** |  |  |  |  |  |  |
| Low, *n* |  | 0 | 0 | 0 | 1 |  |
| Moderate, |  | 2 | 0 | 1 | 3 |  |
| Good, *n* |  | 5 | 6 | 6 | 3 |  |
| Very good, *n* |  | 3 | 3 | 2 | 3 |  |

### Sex-specific CD4 Th cell subset profiles observed in established RA

Percentage of Th1 Th2 Th17 subsets in peripheral blood of participants were determined by flow cytometry **(Figure 2A-2B)**. A pacmap of pooled CD4+CD8-T cells from all participants highlights the clustering of subsets identifying relationships between different Th groups based on surface markers CXCR3 (Th1) CCR4 (Th2), CCR6 (Th17) (**Figure 2A**). We compared Th subset frequencies in RA to control participants and no differences were observed (**Supplementary Figure 2A**). However, when biological sex was included in the stratification of participants, clear differences were observed in Th1, Th17 and Th2 cell frequencies (**Figure 2B**). Th1 cells were lower represented in RA males compared to controls (*p*=0.035), while Th17 cells were higher in males with RA compared to controls (*p=*0.026). Moreover, RA females had lower levels of Th2 cells (*p*=0.023) compared to sex-matched control. Subsequently, RA females had higher Th1 to Th2 cell ratios than controls, while RA males had higher Th17 to Th1 cell ratios than controls. Interestingly, these different Th cell profiles did not correspond to cytokine levels in the blood (**Figure 2C**). In addition, a PCA was performed illuminating participant clustering based on immune parameters (**Figure 3)**. It was notable that participants clustered preferentially by sex rather than RA status implying a relationship between sex and Th cells exists in established RA. This set of data exhibited sex-specific bias toward a Th1TH17-like profile in RA but are not associated with either the progression of the pathology nor with a drastic alteration of cytokine profile.

**Figure 1.**
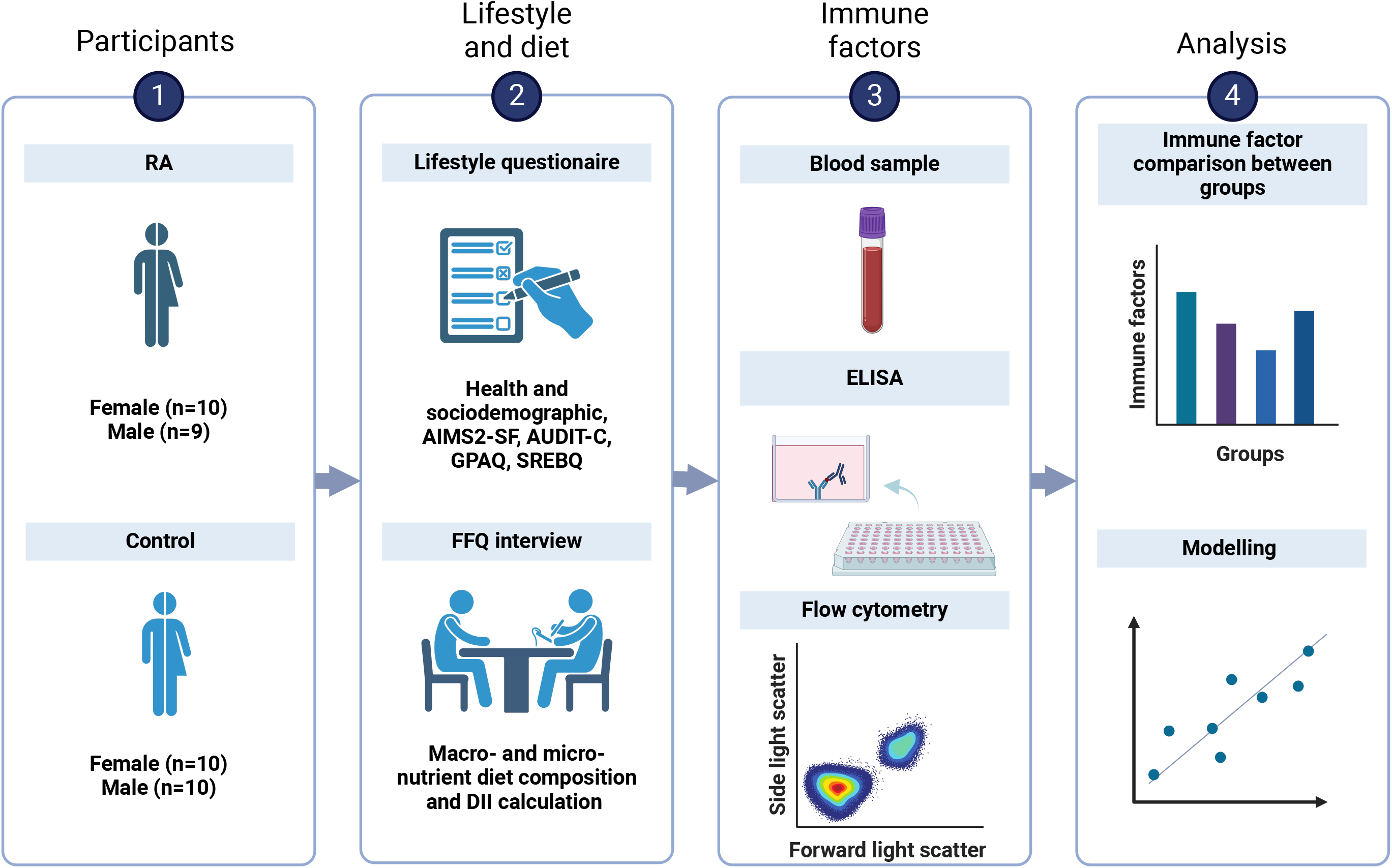
Graphical representation of study participants and overview of experiment design. AIMS2-SF: Arthritis Impact Measurement score short form, AUDIT-C: Alcohol Use Disorders Identification Test, DII: Dietary Inflammatory Index, FFQ: Food Frequency Questionnaire, GPAQ: Global Physical Activity Questionnaire, SHREBQ: Self-Regulation of Eating Behaviour Questionnaire.

**Figure 2.**
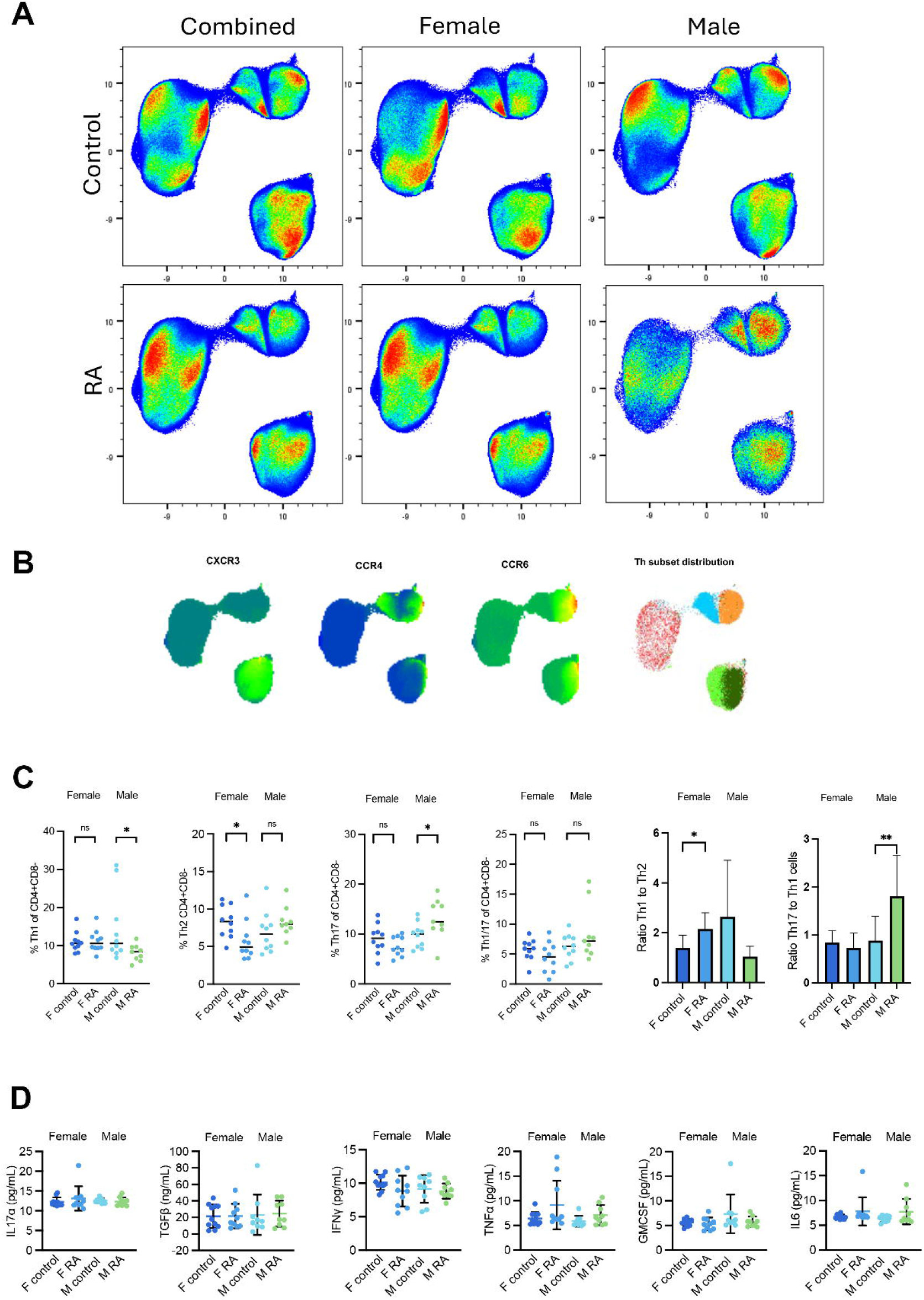
Immune profiles of individuals with RA and healthy controls stratified by sex. (A) Pacmap of CD4 Th cell populations in RA and healthy control groups split by sex, gated on pooled CD4+CD8-cells. (B) Distribution of cell surface marker expression (CXCR3, CCR4 and CCR6). Clustering of Th cells Th1/17 = dark green, Th1 = light green, Th17 = orange, Th2 = blue and other = red. (C) CD4 Th subset frequencies between groups. Horizontal bars indicate the median. Two-tailed Mann-Whitney *U* test *p≤0.05, **p≤0.01, ***p≤0.001. (D) Serum cytokine levels of all participants. Graphs plot individual levels and horizontal bars indicate means ± standard deviation. No significant differences were observed p≤0.05. F = Female, M = Male. RA = Rheumatoid arthritis, control = healthy controls.

### Circulating CD4 Th cells and cytokine levels are predictive of established RA

We next aimed to determine whether there was a relationship between RA diagnosis and combinations of specific circulating immune factors. To this end, we performed a least squares linear regression modelling on immune factors including Th cell frequencies, and cytokine concentrations with an outcome of RA status. Our modelling revealed that individually only IL-6 and IFN-g were significant predictors of RA status (**Supplementary Table 3**). However, the combination of TNFα, IFNγ, IL6, Th1/17 and Th1 cell levels created an 80% model fit and were significantly predictive of RA status (F-statistic 1.05e^-05^) (**Supplementary Table 3**). These data highlight that a combination of biological variables including cytokine levels and Th cell frequencies is a more powerful method to understand RA status than single variables.

### Consumption of dietary components in established RA

Given that diet is an established risk factor in RA, we characterised the mean daily energy and nutrient intakes of the participants through a FFQ interview ^29-30^ **(Table 2)**. Several substantial differences in the estimated consumption of different nutrients were detected between groups. The male RA group consumed 30% less carbohydrate (RA = 156g +/-57g, Control = 253g +/-59g; *p=*0.003), 37% less sugar (RA = 74.1g +/-29g, Control = 118g +/-33g; *p=*0.004) and 30% less dietary fibre (R = 33g +/-8g, Control = 48g +/-13 g; *p=*0.008) than the male control group. The female RA group had the lowest consumption of alcohol in comparison to female control group (RA = 0.31g, Control = 8.97g; *p*=0.047). RA males had the highest DII score in comparison to the RA females with over 4 times the DII (male score: 2.36 ± 1.10, female score: 0.59 ± 1.49, *p<*0.01). Females with RA also had lower levels of vitamin A consumption (RA = 1970μg Control = 2568μg; *p = 0*.*018*). No significant differences were identified for the other 26 food parameters (All *p*’s > 0.05).

**Table 2.** DII scores and daily nutrient intake per group. Different superscript letters within rows indicate statistical difference. DII: Dietary Inflammatory Index. Data presented as mean ± standard deviation or median (1^st^, 3^rd^ quartile). Bolded text indicates significance (p<0.05).

| Dietary components | Female subjects |  | Male subjects |  | P value |
| --- | --- | --- | --- | --- | --- |
|  | RA (n = 10) | Control (n= 10) | RA (n = 9) | Control (n= 10) |  |
| **DII score** | **0.57 ± 1.49<sup>a</sup>** | **0.67 ± 1.31<sup>ac</sup>** | **2.36 ± 1.10<sup>b</sup>** | **2.04 ± 1.55<sup>bc</sup>** | **0.012** |
| **Calorie (kcal)** | 2341 (1930, 2576)<sup>a</sup> | 2334 (2079, 2791)<sup>a</sup> | 2237 (1928, 2715)<sup>a</sup> | 2447 (2174, 2941)<sup>a</sup> | 0.742 |
| **Protein (g)** | 102 (87, 131)<sup>a</sup> | 101 (85, 123)<sup>a</sup> | 121 (87, 170)<sup>a</sup> | 118 (93, 131)<sup>a</sup> | 0.719 |
| **Total fat (g)** | 114 (82, 133)<sup>a</sup> | 94 (69, 116)<sup>a</sup> | 97 (83, 151)<sup>a</sup> | 105 (77, 121)<sup>a</sup> | 0.616 |
| **Saturated fat (g)** | 35.6 ± 9.47<sup>a</sup> | 30.1 ± 13.3<sup>a</sup> | 35.6 ± 11.4<sup>a</sup> | 31.2 ± 9.43<sup>a</sup> | 0.650 |
| **Trans fat (g)** | 1.22 (0.720, 2.08)<sup>a</sup> | 1.12 (0.67, 1.51)<sup>a</sup> | 1.30 (1.07, 3.10)<sup>a</sup> | 1.39 (0.60, 2.00)<sup>a</sup> | 0.613 |
| **Polyunsaturated fat (g)** | 18.4 (13.2, 22.0)<sup>a</sup> | 16.6 (14.0, 18.2)<sup>a</sup> | 14.7 (12.2, 30.0)<sup>a</sup> | 18.0 (14.1, 24.2)<sup>a</sup> | 0.890 |
| **Omega-3 (g)** | 2.33 (2.04, 2.81)<sup>a</sup> | 2.63 (1.40, 3.08)<sup>a</sup> | 2.70 (2.09, 3.92)<sup>a</sup> | 2.68 (1.51, 3.21)<sup>a</sup> | 0.532 |
| **Omega-6 (g)** | 15.3 (10.6, 18.8)<sup>a</sup> | 14.5 (11.3, 15.8)<sup>a</sup> | 10.7 (9.70, 25.6)<sup>a</sup> | 15.1 (11.8, 20.9)<sup>a</sup> | 0.880 |
| **Cholesterol (g)** | 347 ± 115<sup>a</sup> | 280 ± 143<sup>a</sup> | 433 ± 195<sup>a</sup> | 374 ± 135<sup>a</sup> | 0.197 |
| **Carbohydrates (g)** | **201 ± 36.6<sup>ab</sup>** | **232 ± 59.1<sup>ac</sup>** | **156 ± 57.2<sup>b</sup>** | **253 ± 59.8<sup>c</sup>** | **0.003** |
| **Sugar (g)** | **97.8 ± 21.3<sup>ab</sup>** | **118 ± 24.9<sup>a</sup>** | **74.1 ± 29.2<sup>b</sup>** | **118 ± 33.3<sup>a</sup>** | **0.004** |
| **Dietary fibre (g)** | **51.8 ± 14.8<sup>a</sup>** | **53.0 ± 12.1<sup>a</sup>** | **33.6 ± 8.47<sup>b</sup>** | **48.0 ± 13.3<sup>a</sup>** | **0.008** |
| **Alcohol (g)** | **0.31 (0.073, 1.22)<sup>a</sup>** | **8.97 (0.88, 19.7)<sup>b</sup>** | **9.62 (0.22, 14.9)<sup>ab</sup>** | **2.12 (1.08, 7.90)<sup>b</sup>** | **0.047** |
| **Vitamin A (µg)** | **1970 (1667, 2583)<sup>ab</sup>** | **2568 (1788, 3557)<sup>b</sup>** | **1461 (1040, 2069)<sup>a</sup>** | **1436 (953, 2008)<sup>a</sup>** | **0.018** |
| **Thiamin (mg)** | 1.21 (1.09, 2.01)<sup>a</sup> | 1.89 (1.21, 2.31)<sup>a</sup> | 1.49 (1.26, 2.29)<sup>a</sup> | 1.84 (1.47, 2.24)<sup>a</sup> | 0.303 |
| **Riboflavin (mg)** | 1.71 (1.58, 2.30)<sup>a</sup> | 2.31 (1.70, 2.59)<sup>a</sup> | 1.66 (1.44, 2.98)<sup>a</sup> | 2.29 (1.65, 2.96)<sup>a</sup> | 0.523 |
| **Niacin (mg)** | 29.3 (22.9, 40.9)<sup>a</sup> | 30.4 (25.6, 38.7)<sup>a</sup> | 33.8 (28.6, 55.6)<sup>a</sup> | 38.6 (24.9, 50.4)<sup>a</sup> | 0.395 |
| **Vitamin B6 (mg)** | 2.57 ± 0.60<sup>a</sup> | 2.73 ± 0.55<sup>a</sup> | 2.83 ± 0.61<sup>a</sup> | 2.77 ± 1.07<sup>a</sup> | 0.875 |
| **Vitamin B12 (mcg)** | 3.22 (2.58, 4.14)<sup>a</sup> | 3.73 (2.09, 5.53)<sup>a</sup> | 4.30 (3.15, 6.34)<sup>a</sup> | 4.28 (4.05, 6.96)<sup>a</sup> | 0.190 |
| **Vitamin C (mg)** | 217 (166, 258)<sup>a</sup> | 288 (204, 341)<sup>a</sup> | 197 (132, 270)<sup>a</sup> | 236 (131, 318)<sup>a</sup> | 0.223 |
| **Vitamin E (mg)** | 27.6 ± 8.80<sup>a</sup> | 25.4 ± 4.87<sup>a</sup> | 23.5 ± 8.10<sup>a</sup> | 26.6 ± 9.05<sup>a</sup> | 0.710 |
| **Folate (mcg)** | 606 (547, 757)<sup>a</sup> | 728 (615, 935)<sup>a</sup> | 648 (480, 816)<sup>a</sup> | 709 (664, 878)<sup>a</sup> | 0.231 |
| **Sodium (mg)** | 1824 (1667, 2583)<sup>a</sup> | 2492 (1756, 3303)<sup>a</sup> | 2150 (1781, 2974)<sup>a</sup> | 2351 (1696, 4109)<sup>a</sup> | 0.519 |
| **Potassium (mg)** | 4825 (3984, 6335)<sup>a</sup> | 5241 (4791, 6413)<sup>a</sup> | 4481 (4118, 5267)<sup>a</sup> | 4670 (4123, 5892)<sup>a</sup> | 0.485 |
| **Magnesium (mg)** | 568 ± 147<sup>a</sup> | 564 ± 113<sup>a</sup> | 507 ± 125<sup>a</sup> | 566 ± 109<sup>a</sup> | 0.680 |
| **Calcium (mg)** | 969 ± 212<sup>a</sup> | 1179 ± 222<sup>a</sup> | 962 ± 286<sup>a</sup> | 1104 ± 264<sup>a</sup> | 0.185 |
| **Phosphorus (mg)** | 1749 ± 362<sup>a</sup> | 1864 ± 336<sup>a</sup> | 1930 ± 513<sup>a</sup> | 1974 ± 394<sup>a</sup> | 0.633 |
| **Iron (mg)** | 16.7 (13.7, 20.5)<sup>a</sup> | 17.2 (13.5, 20.8)<sup>a</sup> | 12.8 (11.1, 16.5)<sup>a</sup> | 15.8 (14.3, 20.1)<sup>a</sup> | 0.248 |
| **Zinc (mg)** | 12.6 (10.6, 17.3)<sup>a</sup> | 13.4 (10.8, 14.9)<sup>a</sup> | 13.4 (10.5, 16.8)<sup>a</sup> | 14.3 (12.2, 15.3)<sup>a</sup> | 0.799 |
| **Selenium (mg)** | 86.9 (69.6, 108)<sup>a</sup> | 79.4 (60.6, 96.8)<sup>a</sup> | 106 (74.0, 142)<sup>a</sup> | 100 (88.1, 125)<sup>a</sup> | 0.1737 |
| **Iodine (mg)** | 121 ± 26.5<sup>a</sup> | 148 ± 60.7<sup>a</sup> | 133 ± 60.9<sup>a</sup> | 182 ± 50.4<sup>a</sup> | 0.061 |

## Discussion

Sex-based differences in peripheral CD4 Th subsets in established RA have yet to be explored. We observed sex-specific frequencies of Th1, Th2 and Th17 cells in individuals with established RA raising questions regarding differences in RA presentation and management between the sexes. One study has reported sex-based differences in the T cell repertoire of individuals with untreated early RA^34^; here, both male subjects and female subjects with RA had lower Th1/17 frequencies than healthy controls, while females had a higher percentage of Th17s and Th2s, and lower percentage of Th1s than healthy controls. Our findings contrast to this previous study, we observed that RA females had lower percent Th2 cells, and RA males had lower percent Th1 cells and higher percent Th17s than their control counterparts. While these differences are important, it is not surprising to see a contrast between cell profiles in early untreated verses established treated RA, potentially because of exposure to different medications. For example, DMARDs have been shown to alter Th17 frequencies in RA, with decreases observed previously ^35^. In our study, without stratifying by sex, we saw no difference in the frequencies of Th17 cells. After applying the stratification by sex this finding remained for females, while a significant increase in Th17 cells was observed in RA males. Our findings highlight potential immunological differences between males and females with established RA.

Investigation of the predictive nature of CD4 Th cells and associated immune factors in established RA has yet to be described. Our linear regression model highlights TNFα, IFNγ, IL6, Th1/17 and Th1 cells are together predictive of RA in our cohort. Specifically, RA diagnosis in our established RA cohort was associated with enrichment of Th1 cells, TNFα and IL6, and depletion of IFNγ and Th1/17 cells. It is well established that TNFα, IL6 and Th1 cells play a pivotal role in the inflammatory response in RA. Therefore, it was unsurprising that these factors were elevated, even in controlled disease. The depletion of Th1/17 cells and IFNγ in this modelling was also not entirely surprising, likely due to the activity of medication. DMARDs can have different relationship with Th1/17s in different cohorts. For example in juvenile patients, increases in Th1/17 frequencies after treatment have been observed ^36^, while older patients exhibited no change in frequencies after treatment ^37^. Importantly, the participants were overwhelmingly female indicating the potential confounder of sex. Our findings suggest that sex-specific differences in CD4+ T helper subsets could inform more personalized treatment strategies for RA patients. Given that Th1 and Th17 cells are linked to disease activity and severity, sex-based immunological profiling could help clinicians tailor treatment choices, such as the selection of biologics or DMARDs, based on immune phenotypes that are dominant in males and females.

Understanding dietary intake in patients with RA is crucial for studies examining immune cell function. We found that dietary habits varied between males and females living with RA. While our sample size in this pilot study was too low to extrapolate on relationships between dietary consumption patterns and immune cell profiles, this approach is vital to better understand a cohort such as one used in this study. This knowledge can potentially inform targeted nutritional interventions to support disease management.

Despite novelty and research strengths, important limitations of the present study must be declared. We employed an FFQ to broadly determine dietary intake and nutrient composition. However, this technique relies on memory recall, and thus has high variability, is prone to underreporting, and has low accuracy in a small cohort. Therefore, larger cohorts or more sensitive tools that quantify nutrient intake or biomarkers of nutritional status may be beneficial in conducting follow-up studies.

This pilot study proposes a new approach to better understand the relationship between diet and Th cells in RA. From our study, it appears that female and male established RA cohorts could have dimorphic pathophysiology. The levels of Th1, Th2 and Th17 cell frequencies displayed a sex-specific patterns. Several dietary factors including intake of carbohydrates, dietary fibre and alcohol had sex-specific consumption patterns. These results highlight the need for personalized RA management strategies that consider sex-specific immune responses and dietary patterns. Incorporating these findings into clinical practice could enhance therapeutic efficacy and inform tailored dietary recommendations, ultimately improving patient outcomes. Future studies should explore these sex-based dietary-immune interactions in larger cohorts to refine guidelines for personalized RA care.

## Acknowledgements

We thank the participants who volunteered for this study, and Rebecca Davey from Arthritis ACT for her support with participant communication and recruitment. We also acknowledge Harpreet Vohar and Michael Devoy at the flow cytometry facility, John Curtin School of Medical Research, Australian National University, Canberra Australia.

## Funding

This study was supported by a seed grant at the University of Canberra Faculty of Health.

## Disclosure

The authors declare no conflicts of interest.

## Author contributions

SV performed experiments, analysis and wrote the manuscript; CG conceived the project, performed experiments, analysis and edited the manuscript; ND edited the manuscript and conceived the project; TX performed analysis; MK performed experiments; NN, AM, and DP conceived the project; ADR and TT referred patients and provided clinical expertise; JCM, HH and AB provided clinical and/or technical expertise.

## Data availability statement

Data available on request due to privacy/ethical restrictions

## Supplementary Figure and Table legends

**Supplementary Figure 1**. Flow cytometry gating strategy.

**Supplementary Figure 2**. Immune factor comparison between RA and Control groups. (A) Comparison of CD4 Th cell subsets between groups. (B) Comparison of cytokines between groups. Graphs display means with confidence intervals. Comparisons with students T-test. *p≤0.05.

**Supplementary Table 1**. List of Fluorochrome conjugated antibodies used for identification of CD4+ T cell subsets in the present study.

**Supplementary Table 2**. List of medications being taken by RA study participants.

**Supplementary Table 3**. Ordinary least squares (OLS) modelling of immune parameters to predict RA status.

**Supplementary Table 4**. Robust linear modelling identified associations between immune factors and lifestyle variables. Lifestyle variables associated with Th1/17 cells, IL6 and IFNy.

**Supplementary Table 5**. Robust linear modelling of lifestyle variables and immune factors with a sex interaction. Sex-dependant lifestyle variables associated with Th1 cells, IL6 and IFNy.

## Notes

### Competing Interest Statement

The authors have declared no competing interest.

