## Supplementary information for "Characterisation of CD4 Th subsets as well as dietary, exercise and lifestyle factors in an established Rheumatoid Arthritis cohort; pilot study"

Article type: Original Investigation

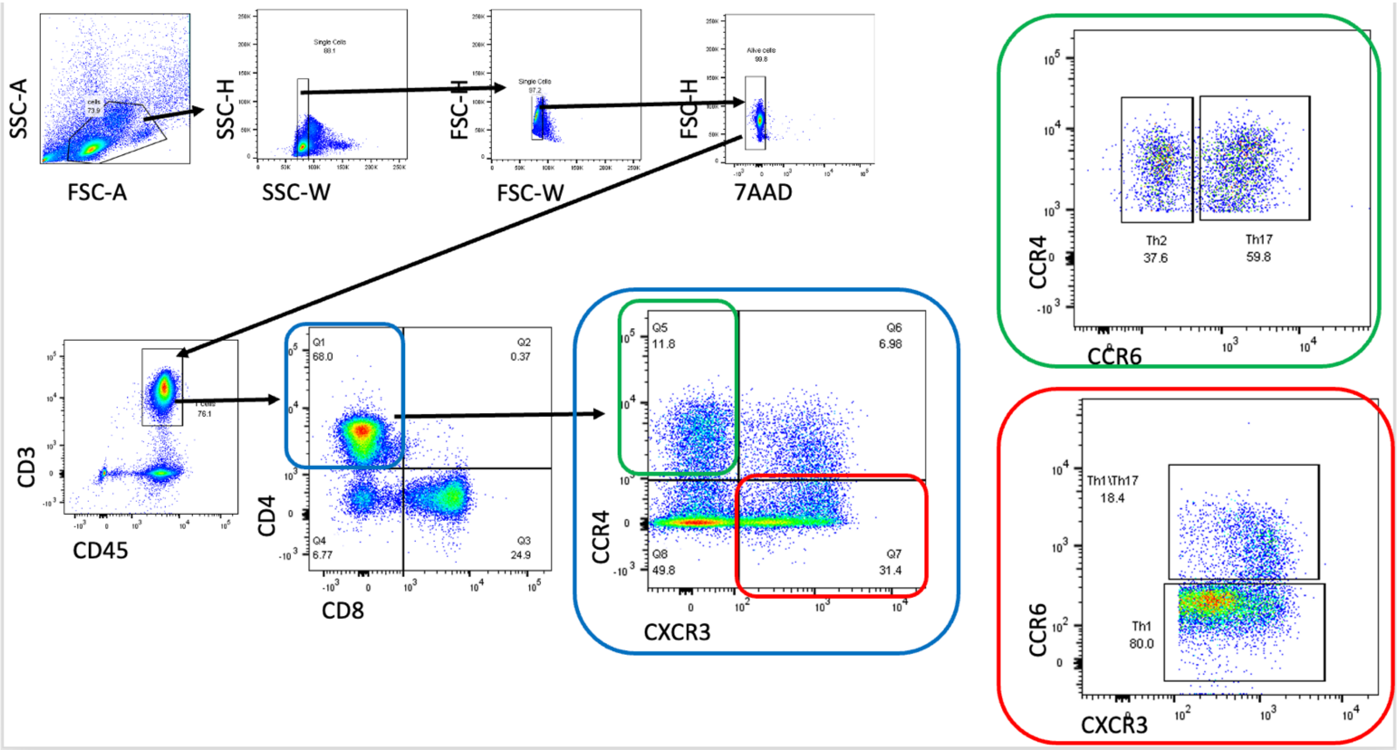

**Supplementary figure 1**. Flow cytometry gating strategy.

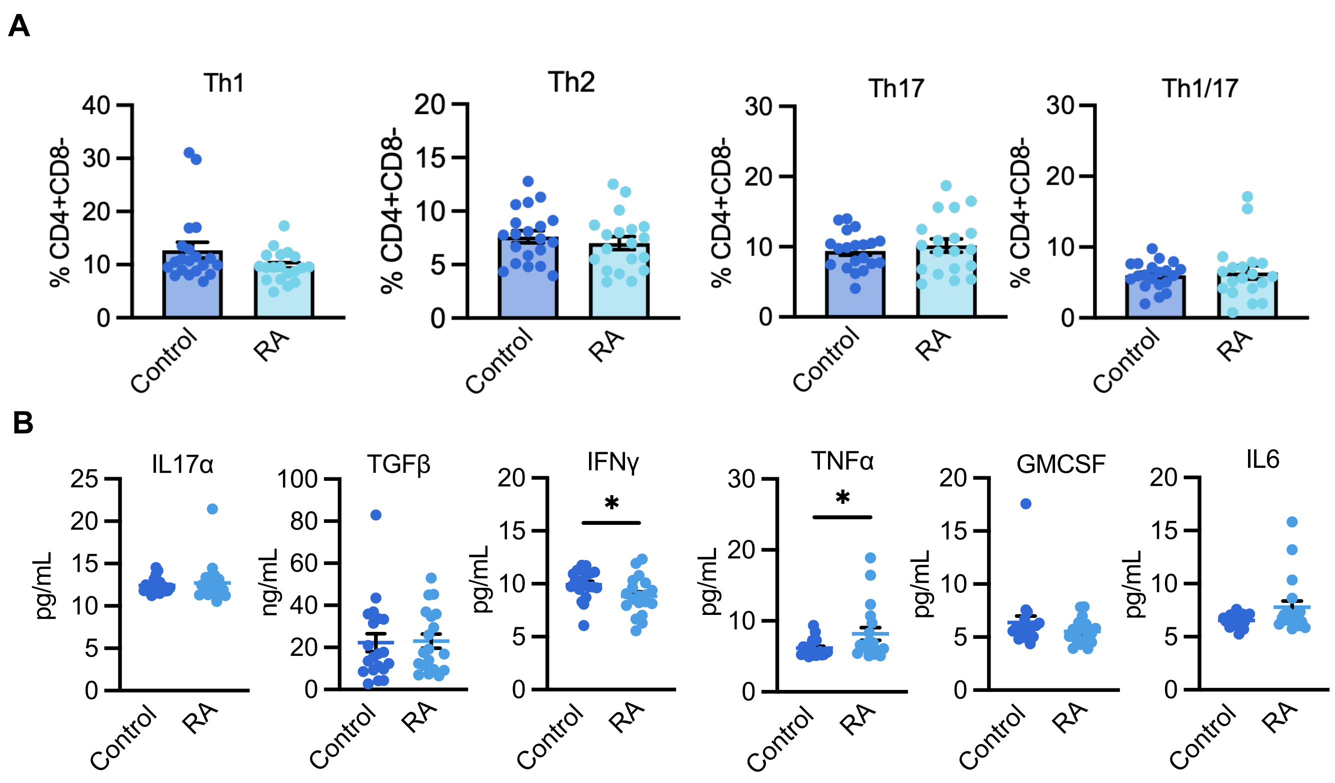

**Supplementary Figure 2.** Immune factor comparison between RA and Control groups. (A) Comparison of CD4 Th cell subsets between groups. (B) Comparison of cytokines between groups. Graphs display means with confidence intervals. Comparisons with students T-test. *p≤0.05.

**Supplementary Table 1.** List of Fluorochrome conjugated antibodies used for identification of CD4+ T cell subsets in the present study.

| Catalog Number | Description | Clone | Isotype |
|---|---|---|---|
| **300316** | PE/Cyanine7 anti-human CD3 | HIT3a | Mouse IgG2a, κ |
| **344714** | APC/Cyanine7 anti-human CD8 | SK1 | Mouse IgG1, κ |
| **317438** | Brilliant Violet 605™ anti-human CD4 | OKT4 | Mouse IgG2b, κ |
| **353208** | Brilliant Violet 421™ anti-human CD197 (CCR7) | G043H7 | Mouse IgG2a, κ |
| **304006** | FITC anti-human CD45 | HI30 | Mouse IgG1, κ |
| **353706** | PE anti-human CD183 (CXCR3) | G025H7 | Mouse IgG1, κ |
| **359408** | APC anti-human CD194 (CCR4) | L291H4 | Mouse IgG1, κ |
| **353408** | Brilliant Violet 421™ anti-human CD196 (CCR6) | G034E3 | Mouse IgG2b, κ |

APC: Allophycocyanin, FITC: Fluorescein isothiocyanate, PE: Phycoerythrin.

**Supplementary Table 2.** List of medications being taken by RA study participants.

| **Medication** | **Female subjects *n*** | **Male subjects *n*** |
|---|---|---|
| Hydroxychloroquine | 2 | 4 |
| Methotrexate | 5 | 3 |
| Sulfasalazine | 1 | |
| Tocilizumab | 1 | 2 |
| Adalimumab | 3 | 1 |
| Upadacitinib | 2 | 1 |
| Tofacitinib | 0 | 1 |
| Certolizumab | 0 | 0 |
| Infliximab | 0 | 0 |
| Etanercept | 0 | 1 |
| **NSAID** | | |
| Meloxicam | 1 | 2 |
| Naprosyn | 0 | 1 |
| Celecoxib | 2 | 2 |
| **Corticosteroids** | | |
| Prednisone | 2 | 3 |

**Supplementary Table 3**. Ordinary least squares (OLS) modelling of immune parameters to predict RA status.

| *Immune parameter* | *coefficient* | *p value* | *Model fit* | *F statistic* |
|---|---|---|---|---|
| *Th1 cells* | 0.0190 | 0.206 | | |
| *TNFa* | 0.0504 | 0.041 | | |
| *Th1/17 cells* | -010185 | 0.111 | | |
| *IL6* | 0.0734 | 0.055 | | |
| *IFNg* | -0.0458 | 0.144 | | |
| *OLS* | | | 80% | $1.05e^{-05}$ |

**Supplementary Table 4.** Robust linear modelling identified associations between immune factors and lifestyle variables. Lifestyle variables associated with Th1/17 cells, IL6 and IFNy.

| | Var name | P val | Coef | Std err | Adjusted P |
|---|---|---|---|---|---|
| Th1/17 cells | Carbohydrate (g) | 0.001897 | 50.917935 | 16.394283 | 0.026835 |
| | Starch (g) | 0.000022 | 47.904942 | 11.282221 | 0.002154 |
| | Vitamin C (mg) | 0.001679 | 34.531042 | 10.99056 | 0.026835 |
| | Total folate (µg) | 0.000631 | 54.238131 | 15.868421 | 0.020233 |
| | Folate,total DFE (µg) | 0.002666 | 58.156465 | 19.360679 | 0.029325 |
| | Food Folate (µg) | 0.004408 | 44.575521 | 15.654802 | 0.033567 |
| | kJ from carbohydrate (%) | 0.000818 | 54.979868 | 16.427593 | 0.020233 |
| | Fat as mono (%) | 0.006272 | 88.690858 | 32.449318 | 0.041395 |
| | GRAINS (serve) | 0.000278 | 36.141342 | 9.94171 | 0.013743 |
| | Refined (serve) | 0.002529 | 28.278858 | 9.364256 | 0.029325 |
| IL6 | sat | 0 | 16.369828 | 2.495183 | 0 |
| | Protein (g) | 0.014135 | 10.183905 | 4.150245 | 0.025367 |
| | Total fat (g) | 0.000882 | 9.039718 | 2.718268 | 0.00371 |
| | Saturated fat (g) | 0.000151 | 9.824644 | 2.592428 | 0.001098 |
| | Trans Fatty Acids (g) | 0.002592 | 9.056027 | 3.006283 | 0.008248 |
| | Monounsaturated fat (g) | 0.007044 | 7.264193 | 2.695689 | 0.015409 |
| | Cholesterol (mg) | 0.013228 | 6.129745 | 2.474098 | 0.024368 |
| | Carbohydrate (g) | 0.000002 | 11.258252 | 2.378081 | 0.000038 |
| | Sugars (g) | 0.000089 | 10.011663 | 2.555206 | 0.000781 |
| | Free Sugars (g) | 0.022791 | 4.972918 | 2.184058 | 0.035109 |
| | Starch (g) | 0.001452 | 7.856149 | 2.467244 | 0.005348 |
| | Water (g) | 0.001003 | 8.434731 | 2.564048 | 0.003902 |
| | Dietary fibre (g) | 0.004149 | 9.512984 | 3.318567 | 0.010373 |
| | Ash (g) | 0.000901 | 11.545871 | 3.477958 | 0.00371 |
| | Niacin (mg) | 0.029871 | 6.419241 | 2.955737 | 0.042674 |
| | Niacin equivalents (mg) | 0.036765 | 7.305262 | 3.498085 | 0.049492 |
| | Vitamin E (mg) | 0.000029 | 13.587566 | 3.246707 | 0.000333 |
| | Tocopherol, alpha (mg) | 0.004308 | 9.523936 | 3.336263 | 0.010399 |
| | Vitamin B6 (by analysis) (mg) | 0.000605 | 11.384451 | 3.319765 | 0.003026 |
| | Food Folate (µg) | 0.010054 | 7.894878 | 3.067205 | 0.021304 |
| | Total vitamin A equivalents (µg) | 0.016473 | 7.01511 | 2.925075 | 0.027455 |
| | Beta carotene equivalents (µg) | 0.018131 | 6.156429 | 2.60542 | 0.028845 |
| | Beta carotene (µg) | 0.014496 | 6.123906 | 2.504931 | 0.025367 |
| | Sodium (mg) | 0.000479 | 7.49609 | 2.146601 | 0.002581 |
| | Potassium (mg) | 0.032747 | 9.863458 | 4.619512 | 0.044947 |
| | Magnesium (mg) | 0.004955 | 11.016873 | 3.920664 | 0.01133 |
| | Calcium (mg) | 0.000175 | 12.776505 | 3.405223 | 0.001116 |
| | Phosphorus (mg) | 0.000157 | 13.715089 | 3.628435 | 0.001098 |
| | Iron (mg) | 0.000712 | 14.204708 | 4.196391 | 0.003322 |
| | Zinc (mg) | 0.003106 | 11.76723 | 3.979424 | 0.008698 |
| | kJ from fat (%) | 0.003074 | 13.6373 | 4.606863 | 0.008698 |
| | kJ from saturated fat (%) | 0.001907 | 12.485659 | 4.021928 | 0.006673 |
| | kJ from carbohydrate (%) | 0.003847 | 9.465572 | 3.27477 | 0.009981 |
| | Fat as mono (%) | 0 | 25.43226 | 4.247482 | 0 |

| | | | | | |
|---|---|---|---|---|---|
| | Fat as poly (%) | 0.025916 | 8.499447 | 3.815755 | 0.038599 |
| | Fat as saturated (%) | 0.000315 | 17.514851 | 4.861778 | 0.001838 |
| | F18D2CN6 linoleic (g) | 0.023072 | 7.925079 | 3.487774 | 0.035109 |
| | F18D3N3 alpha-linolenic (ALA) (g) | 0.000038 | 11.473041 | 2.785274 | 0.00038 |
| | GRAINS (serve) | 0.010631 | 5.623587 | 2.20136 | 0.021304 |
| | Refined (serve) | 0.00385 | 4.98354 | 1.724274 | 0.009981 |
| | Red & orange vegetables (serve) | 0.029713 | 5.010556 | 2.304876 | 0.042674 |
| | Eggs (serve) | 0.002929 | 6.455226 | 2.169774 | 0.008698 |
| | Soy products (serve) | 0.010652 | 4.494984 | 1.760038 | 0.021304 |
| | OIL EQUIVALENTS (tsp) | 0.005017 | 7.358589 | 2.622526 | 0.01133 |
| | UNCLASSIFIED WEIGHT (g) | 0.000007 | 26.484316 | 5.903178 | 0.000101 |
| | · Unclassified weight percent (%) | 0 | 29.399443 | 3.369413 | 0 |
| | Omega 3 | 0.012599 | 7.36342 | 2.951371 | 0.024368 |
| | Alcohol_score | 0.002457 | 12.225988 | 4.036898 | 0.008191 |
| | Physical_activity | 0.015266 | 4.846878 | 1.997888 | 0.026064 |
| | sat | 0.000669 | 29.146264 | 8.566944 | 0.009073 |
| | Carbohydrate (g) | 0.007543 | 18.888682 | 7.069465 | 0.037715 |
| | Sugars (g) | 0.001388 | 22.819716 | 7.137542 | 0.014651 |
| | Dietary fibre (g) | 0.000775 | 22.531245 | 6.702414 | 0.0092 |
| | Ash (g) | 0.000164 | 27.379533 | 7.263402 | 0.003885 |
| | Thiamin (mg) | 0.004837 | 14.112396 | 5.00855 | 0.027033 |
| | Niacin (mg) | 0.006027 | 17.763725 | 6.468176 | 0.031807 |
| | Tocopherol, alpha (mg) | 0.003283 | 21.133859 | 7.188487 | 0.022274 |
| | Vitamin B6 (by analysis) (mg) | 0.01185 | 19.885594 | 7.901814 | 0.046905 |
| **IFNy** | Vitamin B12 (µg) | 0.000052 | -228.65778 | 56.516742 | 0.001651 |
| | Food Folate (µg) | 0.003775 | 19.003905 | 6.561244 | 0.023731 |
| | Sodium (mg) | 0.000635 | 17.006951 | 4.978369 | 0.009073 |
| | Potassium (mg) | 0.002089 | 28.243678 | 9.178214 | 0.018766 |
| | Calcium (mg) | 0.002244 | 25.151274 | 8.230558 | 0.018766 |
| | Iron (mg) | 0.002638 | 29.901084 | 9.943657 | 0.019277 |
| | kJ from fibre (%) | 0.009184 | 21.003473 | 8.062405 | 0.043626 |
| | Fat as mono (%) | 0 | 61.752918 | 10.539506 | 0 |
| | Organ meats (serve) | 0.000028 | -4.605982 | 1.099115 | 0.001321 |
| | - Seafood high in LC N-3 (serve) | 0.011695 | -8.130501 | 3.224851 | 0.046905 |

**Supplementary Table 5.** Robust linear modelling of lifestyle variables and immune factors with a sex interaction. Sex-dependant lifestyle variables associated with Th1 cells, IL6 and IFNy.

| | Var name | P val | Coef | Std err | Adjusted P |
|---|---|---|---|---|---|
| | Cholesterol (mg) | 0.000475 | 38.067167 | 10.893683 | 0.026605 |
| **Th1 cells** | kJ from carbohydrate (%) | 0.000332 | -70.676306 | 19.694009 | 0.026605 |
| | Iron (mg) | 0.001028 | 14.101299 | 4.295605 | 0.038039 |
| **IL6** | | | | | |
| | Vitamin B12 (µg) | 0.000049 | 228.673257 | 56.348279 | 0.002744 |
| | Retinol (µg) | 0.000238 | 22.707994 | 6.178978 | 0.008799 |
| **IFNy** | Organ meats (serve) | 0.000028 | 4.605982 | 1.099115 | 0.002744 |
